# Not enough pleasure? Influence of hallucination proneness on sensory feedback processing of positive self-voice

**DOI:** 10.1101/2024.12.04.626770

**Authors:** Suvarnalata Xanthate Duggirala, Michael Schwartze, Lisa K. Goller, David E. J. Linden, Ana P. Pinheiro, Sonja A. Kotz

**Affiliations:** Department of Neuropsychology and Psychopharmacology, Faculty of Psychology and Neuroscience, Maastricht University, Maastricht, Netherlands; Faculty of Psychology, University of Lisbon, Lisbon, Portugal; Department of Psychiatry and Neuropsychology, School for Mental Health and Neuroscience, Faculty of Health and Medical Sciences, Maastricht University, Maastricht, Netherlands; Maastricht University Medical Center, Maastricht, Netherlands

**Keywords:** N100-P200-N200, motor-auditory task, pleasure, certainty, hallucination proneness

## Abstract

Ample research explored changes in sensory feedback processing of the self-voice as well as the control of attention allocation in voice hearers, including both non-clinical voice hearers and voice hearers with a psychotic disorder. While the attentional bias toward negative emotional information in voice hearers with a psychotic disorder due to heightened sensitivity towards threat/danger is well established, it remains unclear how positive emotion captures or controls attention. Manipulating the certainty of sensory feedback to the self-voice, transitioning from fully neutral to entirely positive (100% neutral, 60-40% neutral-pleasure, 50-50% neutral-pleasure, 40-60% neutral-pleasure, 100% pleasure), provides an opportunity to investigate attentional control and sensory feedback processing in positive self-voice as a function of hallucination proneness (HP). Participants with different HP scores self-generated and passively listened to their own voices during EEG recordings. N100 or P200 responses to self-generated and externally-generated self-voices did not differ. Further, HP did not modulate N100/P200 responses. These null findings might result from the minimal perceptual discriminability among the five types of voices varying in pleasure content. This might have led to less variation in certainty regarding the sensory feedback to self-voice, and consequently a lack of differential engagement of attentional resources. The lack of a global N100 suppression effect prompts inquiry into the association of sense of ownership/agency and pleasure.

## 1. Introduction

Although typically associated with a clinical diagnosis of psychiatric (e.g., schizophrenia, bipolar disorder, or major depressive disorder) or neurological (e.g., Alzheimer’s disease, Parkinson’s disease) disorder, auditory verbal hallucinations (AVH) are also present in healthy individuals not in need of clinical care (Daalman et al., 2011; Johns, Hemsley, & Kuipers, 2002; Johns et al., 2014; Larøi, 2012; Maijer, Begemann, Palmen, Leucht, & Sommer, 2018; van Os, 2003). The differences between psychotic and non-clinical AVH pertain to their emotional valence, controllability, and related distress (Daalman et al., 2011; Johns et al., 2002; Johns et al., 2014; Larøi, 2012; Maijer et al., 2018). A growing body of evidence suggests that the neural underpinnings (Allen, Aleman, & McGuire, 2007; Barkus, Stirling, Hopkins, McKie, & Lewis, 2007; Brebion et al., 2016; Diederen et al., 2012) and the experience of AVH in terms of characteristics such as sound location, loudness, and source are similar in psychotic and non-clinical voice hearers (Daalman et al., 2011; Johns et al., 2002; Johns et al., 2014; Larøi, 2012; Maijer et al., 2018). This evidence and the prevalence rates of AVH may reflect a continuum of susceptibility in the general population ranging from low to high hallucination proneness (HP; (Baumeister, Sedgwick, Howes, & Peters, 2017; Johns et al., 2002; van Os, Linscott, Myin-Germeys, Delespaul, & Krabbendam, 2009).

One of the theories embedded in the ‘forward model’ framework that accounts for AVH relates to self-monitoring and inner-speech (Feinberg, 1978; Frith, Friston, Liddle, & Frackowiak, 1992; Frith, Blakemore, & Wolpert, 2000; Jones & Fernyhough, 2007). According to this theory, alterations in self-monitoring may lead to the attribution of one’s own actions to an outside agent. This means that if AVH are a form of inner speech, voice hearers may fail to recognize them as self-generated. The forward model suggests that self-compared to externally-generated actions and their sensory consequences lead to a suppression of neural activity as individuals can fully predict the sensory consequences of their own actions (Blakemore, Wolpert, & Frith, 2000; Frith et al., 2000; Wolpert, Ghahramani, & Jordan, 1995). Any disruption of this internal ‘self-monitoring’ system may result in the misattribution of internally-generated speech to an external source, and consequently lead to the manifestation of experiences such as AVH. Electroencephalographic (EEG) studies in voice hearers provide support for this conception (Ford & Mathalon, 2004; Ford, Mathalon, Heinks, et al., 2001; Ford, Mathalon, et al., 2001a, 2001b; Ford et al., 2013; Pinheiro et al., 2020; Pinheiro, Schwartze, & Kotz, 2018). Voice hearers with a psychotic disorder and non-clinical voice hearers both showed altered processing of self-generated voices, as indicated by an increased N100 event-related potential (ERP) response to self-compared to externally-generated voices (Ford & Mathalon, 2004; Ford, Mathalon, Heinks, et al., 2001; Ford, Mathalon, et al., 2001a, 2001b; Ford et al., 2013; Pinheiro et al., 2020; Pinheiro et al., 2018). Alternatively, it might indicate error in attention allocation and control whereby voice hearers allocate attention to a contextually irrelevant stimulus (e.g., self-generated own voice). Given that these alterations are observed in both voice hearers with and without a diagnosis of a psychotic disorder, they potentially support the psychosis continuum hypothesis.

Emotions provide cues that can crucially influence the memory of an event in the source monitoring framework (Johnson, Hashtroudi, & Lindsay, 1993). Emotional quality of the hallucinated voices not only offers contextual information about the event’s source but also imparts details about the distinctiveness of a voice. Any alteration in processing emotional characteristics may therefore increase the risk of misattribution of internally-generated sensations to external sources. The emotional quality of hallucinated voices is one of their main characteristics. They are more often negative and derogatory for voice hearers with a psychotic disorder and more positive and neutral for non-clinical voice hearers (Daalman et al., 2011; Johns et al., 2014; Larøi, 2012). This difference in the emotional quality of AVH in non-clinical voice hearers and voice hearers with a psychotic disorder has also been reported in vocal emotion processing (Amminger, Schafer, Klier, et al., 2012; Amminger, Schafer, Papageorgiou, et al., 2012; Amorim, Roberto, Kotz, & Pinheiro, 2022; van ‘t Wout, Aleman, Kessels, Laroi, & Kahn, 2004). Voice hearers with a psychotic disorder tend to perceive negative emotions more intensely and to misattribute negative meaning to a neutral stimulus, potentially due to altered predictions causing them to constantly anticipate negative sensations and pay involuntary attention to negative cues (Mohanty et al., 2008). Further, voice hearers with a psychotic disorder tend to show low positive affect, reduced pleasure expression and recognition (Cohen & Minor, 2010; Horan, Blanchard, Clark, & Green, 2008; Kring & Moran, 2008; Li, Fung, Moore, & Martin, 2019; Watson & Naragon-Gainey, 2010). This reduced ability to process positive emotions is closely associated with negative symptoms such as social aloofness and constricted affect (Watson & Naragon-Gainey, 2010). If the heightened perception of negative emotions and the tendency to attribute negative connotations to neutral stimuli among voice hearers with a psychotic disorder are linked to their inherent attentional bias favoring negative cues (Alba-Ferrara, de Erausquin, Hirnstein, Weis, & Hausmann, 2013; Birchwood & Chadwick, 1997; Galdos et al., 2011; Nelson, Whitford, Lavoie, & Sass, 2014a, 2014b; Rossell & Boundy, 2005), it can be hypothesized that non-clinical voice hearers, who frequently encounter positive and neutral voices, may have an enhanced ability to discern positive content within neutral stimuli. By manipulating positive emotional quality it may be possible to vary certainty about sensory feedback to self-voice production as well as probe control of attention allocation. This manipulation may allow disentangling sensory feedback processing and control of attention allocation in non-clinical individuals who are highly prone to hallucinations. Understanding these processes within the context of positive emotion as a function of HP in non-clinical individuals, might help develop interventions to improve diminished pleasure perception and motivation in voice hearing (Foussias, Agid, Fervaha, & Remington, 2014; Nguyen et al., 2016).

The excellent temporal resolution of EEG allows sensitive monitoring of dynamical changes in voice quality. Here, we extended prior work (Pinheiro et al., 2020; Pinheiro et al., 2018), using a well-validated EEG motor-auditory task (figure 1) where the self-voice changes from fully neutral to fully pleasure-100% neutral, 60-40% neutral-pleasure; 50-50% neutral-pleasure; 40-60% neutral-pleasure and 100% pleasure. We focused on the N100, P200, and N200 ERP responses as established indicators of sensory feedback processing, conscious differentiation between self-generated and externally-generated events, and attention allocation and error awareness. For 100% neutral and 100% pleasure self-voice, we expected to observe reduced N100 and P200 suppression effects (self-> externally-generated) with increased HP. The hypotheses for the direction of the effect of uncertain/ambiguous self-voices were more exploratory. Previous studies (Addington, Penn, Woods, Addington, & Perkins, 2008; Amminger, Schafer, Klier, et al., 2012; Amminger, Schafer, Papageorgiou, et al., 2012; Amorim et al., 2022) showed that participants scoring high on HP tend to miscategorize vocal emotions, in particular, ambiguous voices from the neutral-pleasure continuum were categorized as ‘angry’. Therefore, we expected an increase in N100, P200 as well as N200 amplitude for self-than externally-generated ambiguous (uncertain: 60-40%: neutral-pleasure; 50-50%: neutral-pleasure; 40-60%: neutral-pleasure) self-voices in individuals with low compared to high HP individuals. The underlying rationale was that individuals characterized by low HP would display changes in both sensory feedback processing and error perception and awareness when listening to unexpected and ambiguous self-voices.

**Figure 1:**
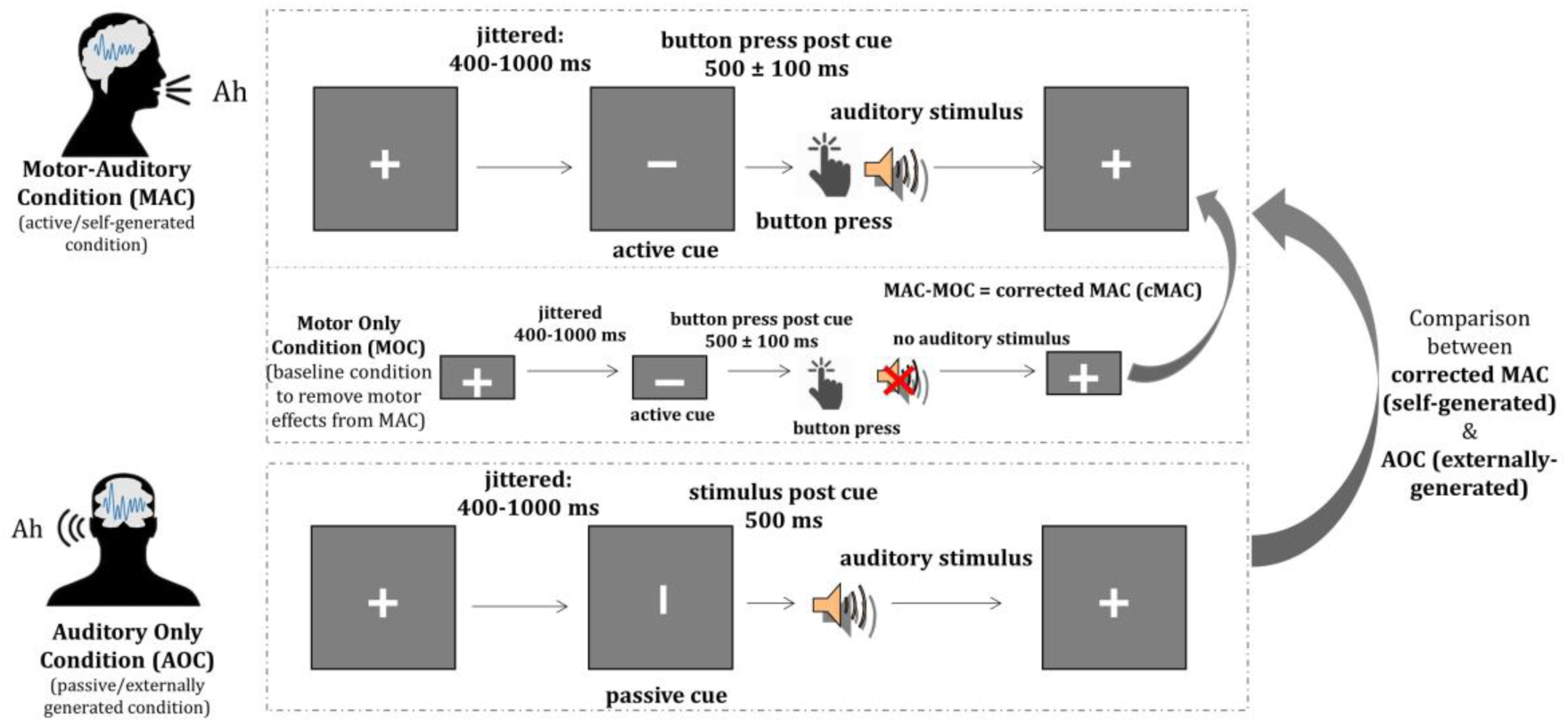
Graphical representation of the Motor-auditory task. Abbreviations: MA = Motor Auditory Condition; AO = Auditory Only Condition; MO = Motor Only Condition. Motor activity from MA condition was removed by subtracting MO from MA to obtain MA corrected condition. Statistical analyses were performed with ERPs from MAc and AO conditions.

## 2. Methods

### Participants

Twenty-nine healthy adults (age range 18-27 years) were recruited. All participants were first invited for a voice recording, followed by the EEG session. Three participants did not participate in the EEG sessions due to time constraints, whereas one participant was excluded from further analysis due to technical issues during the EEG data collection. Therefore, the final participant number was 25 (21 females, mean age = 21.24, s.d. = 2.49 years; 21 right-, 3 left-handed, and 1 ambidextrous) varying in HP (Launay Slade Hallucination Scale (LSHS)(Castiajo & Pinheiro, 2017; Larøi & Van der Linden, 2005a, 2005b; Launay & Slade, 1981) total scores: mean = 18.56, s.d. = 10.17, max = 42, min = 3; LSHS AVH scores [sum of items: “In the past, I have had the experience of hearing a person’s voice and then found no one was there”, “I often hear a voice speaking my thoughts aloud”, and “I have been troubled by voices in my head”]: mean = 2.40, s.d. = 2.62, min = 0, max = 11). All participants provided their written informed consent before the start of the study. They either received financial compensation (vouchers) or study credits for their participation. All participants self-reported normal or corrected-to-normal visual acuity and normal hearing. The study was approved by the Ethics Committee of the Faculty of Psychology and Neuroscience at Maastricht University and performed in accordance with the approved guidelines and the Declaration of Helsinki (ERCPN-176_08_02_2017_S2).

### Procedure

All participants went through the following sessions performed in two separate visits. Session 1: Voice recording and stimulus generation

### Voice recording

Participants comfortably sat inside an acoustically and electrically shielded chamber with the recording equipment, while the researcher sat outside this chamber. Recordings were made using a Rode NTKb microphone powered by a Rode NTK microphone power supply (http://www.rode.com/microphones/ntk) and processed with the Praat software (https://www.praat.org). Participants were instructed to repeatedly vocalize “ah” and “oh” in a neutral (no emotion) and in a pleasure voice. Vowels were chosen to eliminate semantic content (Cook & Wilding, 2001; Schweinberger, Herholz, & Sommer, 1997; Ventura, Nagarajan, & Houde, 2009). Participants were asked to vocalize the vowels for 500 ms, and were provided with examples to familiarize them with the target duration of the vocalization. This duration was chosen to properly capture the emotionality while maintaining adequate length for self-voice recognition. The best voice samples were selected once the participants confirmed that they recognized their recorded voice, that the pleasure intensity was the highest that they could produce, that they perceived no emotion in the neutral recording, and if the vowels were pronounced clearly. Background noise was eliminated from the recordings using Audacity software (https://audacityteam.org/) and a Praat script was applied to normalize the intensity to 70 dB. The duration of the final neutral and pleasure “ah” and “oh” vocalizations for each participant was 500 ms.

### Morphing

To create voice samples with varying degrees of emotional content, the pre-recorded neutral and pleasure self-voices for each individual participant were parametrically morphed to create neutral-to-pleasure and pleasure-to-neutral continua. These continua consisted of 11 stimuli with a 10% stepwise increase (neutral-to-pleasure)/decrease (pleasure-to-neutral) in emotionality along the continuum (see supplementary table 1). Morphing was performed using the TANDEM-STRAIGHT software (Kawahara, 2006; Kawahara & Irino, 2005; Kawahara et al., 2008) running on MATLAB (R2019a, v9.6.0.1072779, The MathWorks, Inc., Natick, MA). For the final EEG experiments, 100% neutral, 60-40%: neutral-pleasure; 50-50%: neutral-pleasure; 40-60%: neutral-pleasure and 100% pleasure voice morphs were selected. The intermediate voice morphs were selected based on pilot data that revealed that the maximum uncertainty to differentiate a neutral from a pleasure self-voice fell in the range of 35-65% morphing. The increase in emotional voice quality (as self-voice stepwise changes from fully neutral to fully pleasure) and manipulations of uncertainty (most certain: 100% neutral and pleasure; uncertain: 60-40%: neutral-pleasure; 50-50%: neutral-pleasure; 40-60%: neutral-pleasure self-voice morphs) would probe both changes in sensory feedback to the self-voice and attention allocation resulting from these changes.

#### Session 2: EEG

Participants were given an overview of the procedure and the principles of EEG at the start of the session. They sat comfortably in an electrically shielded soundproof chamber in front of a screen placed about 100 cm away. Participants filled in the LSHS questionnaire while the EEG cap was prepared.

### Auditory-motor task

A variant of an established button-press task was employed to investigate differences between responses to self-and externally-generated auditory stimuli (Pinheiro et al., 2020; Pinheiro et al., 2018) (figure 1). This task comprises three conditions: a motor-auditory condition (MAC), where participants pressed a button to generate their pre-recorded voice; an auditory-only condition (AOC), where participants passively listened to their pre-recorded voice; and a motor-only condition (MOC), where they pressed a button but did not hear their voice. This latter condition was used to control for motor activity resulting from the button-press in the MA condition (MAC-MOC = corrected MAC [cMAC]). Previous studies have consistently shown that there is a reduction in the N100 amplitude in response to self-generated sound via a button-press compared to passively listening to the same sound (Baess, Widmann, Roye, Schroger, & Jacobsen, 2009; Hughes, Desantis, & Waszak, 2013), indicating that button-presses can be used as a motor-act to approximate self-generation of a speech stimulus (for voices see (Knolle, Schwartze, Schroger, & Kotz, 2019).

The paradigm was presented in a fully randomized event-related design over 12 runs. Each run consisted of 80 trials (40 AO, 40 MA, and 10 MO). Each trial started with a fixation cross, after which the presentation (vertical or horizontal) of a cue was jittered between 400-1000 ms. The cue was then followed by an auditory stimulus (after 500 ms for AO) or a button press that may (MA) or may not (MO) elicit an auditory stimulus. Five types of voice morphs consisting of “ah” and “oh” vocalizations, respectively, were presented in the AO and MA conditions. Thus, each run consisted of 4 trials of 10 stimulus types each (“ah” and “oh” for 5 voice morphs). This included 96 trials per voice morph (“ah” and “oh” combined, supplementary table 1). Participants were given short breaks after each run. To minimize potential influences of lateralized motor activity, participants were asked to switch their response hand every three runs. Prior to the experiment, participants were trained to press the button within 500 ± 100 ms after the cue (horizontal bar) to align the presentation of auditory stimuli in the MA and AO conditions and to avoid overlap of cue-elicited and motor activation.

The task was programmed and presented using the Presentation software (version 18.3; Neurobehavioral Systems, Inc.). Auditory stimuli were presented via in ear inserts. Button presses were recorded via the spacebar button on the keyboard.

### Stimulus Rating

At the end of the EEG session, participants rated their voices for arousal and valence (supplementary figure 1). They additionally rated the voices on perceived ownness, i.e., how much they identified their self-voice on a Likert scale (0-10). This was done to ensure that participants recognized their own voice and perceived the emotion expressed by it.

### EEG data acquisition and preprocessing

EEG data were recorded with BrainVision Recorder (Brain Products, Munich, Germany) using an ActiChamp 128-channel active electrode setup while participants performed the auditory-motor task. Data were acquired with a sampling frequency of 1000 Hz, an electrode impedance below 10 kΩ, using TP10 as online reference. EEG data were pre-processed using the Letswave6 toolbox (https://github.com/NOCIONS/letswave6) running on MATLAB 2019a. Data were first cleaned to remove false button presses (e.g., trials with button presses during AO), downsampled to 500 Hz, and then bandpass filtered (1-30 Hz). All channels were re-referenced to the average of the mastoid electrodes. Eye blinks and movements and noisy electrodes were removed using an independent component analysis (ICA) with the runica algorithm in combination with Rajan and Rayner (PICA) as implemented in Letswave6 (https://github.com/NOCIONS/letswave6). ICs representing primarily noise were removed for each participant based on the IC time course and topography. The resulting data were segmented using a-600 to 800 ms time-window relative to the onset of the auditory stimulus. The segmented data were baseline corrected to a-600 to-400 ms window. This remote baseline was selected due to a cue-related ERP modulation before the onset of the auditory stimulus in AO. After baseline correction, an automatic artifact rejection algorithm was applied with an amplitude criterion of ± 65µV to remove epochs/trials with remaining artifacts. The resulting data were then averaged for each participant and each condition. The grand averaged waveforms revealed three ERP components, two negative components peaking at 164 ms and 460 ms and one positive component peaking at 286 ms. As the latencies of the ERP responses varied significantly (supplementary table 2), peak amplitudes as an outcome measure were chosen for data quantification. The N100 peak amplitude was defined as the largest negative peak occurring between 80-230 ms, the P200 peak amplitude was defined as the following positive peak between N100 and 380 ms, and the N200 peak amplitude as the negative peak between the P200 and 600 ms (Swink & Stuart, 2012a, 2012b). Previous research showed that the ERP components of interest all have prominent fronto-medial and fronto-central topographies (Behroozmand, Karvelis, Liu, & Larson, 2009; Chen, Chen, Liu, Huang, & Liu, 2012; Korzyukov, Karvelis, Behroozmand, & Larson, 2012). Therefore, the N100, P200, and N200 responses were extracted from the same fronto-central region of interest (ROI) that included 21 electrode locations: AFF1h, AFF2h, F1, Fz, F2, FFC3h, FFC1h, FFC2h, FFC4h, FC3, FC1, FCz, FC2, FC4, FCC3h, FCC1h, FCC2h, FCC4h, C1, Cz, C2 (figure 2).

**Figure 2:**
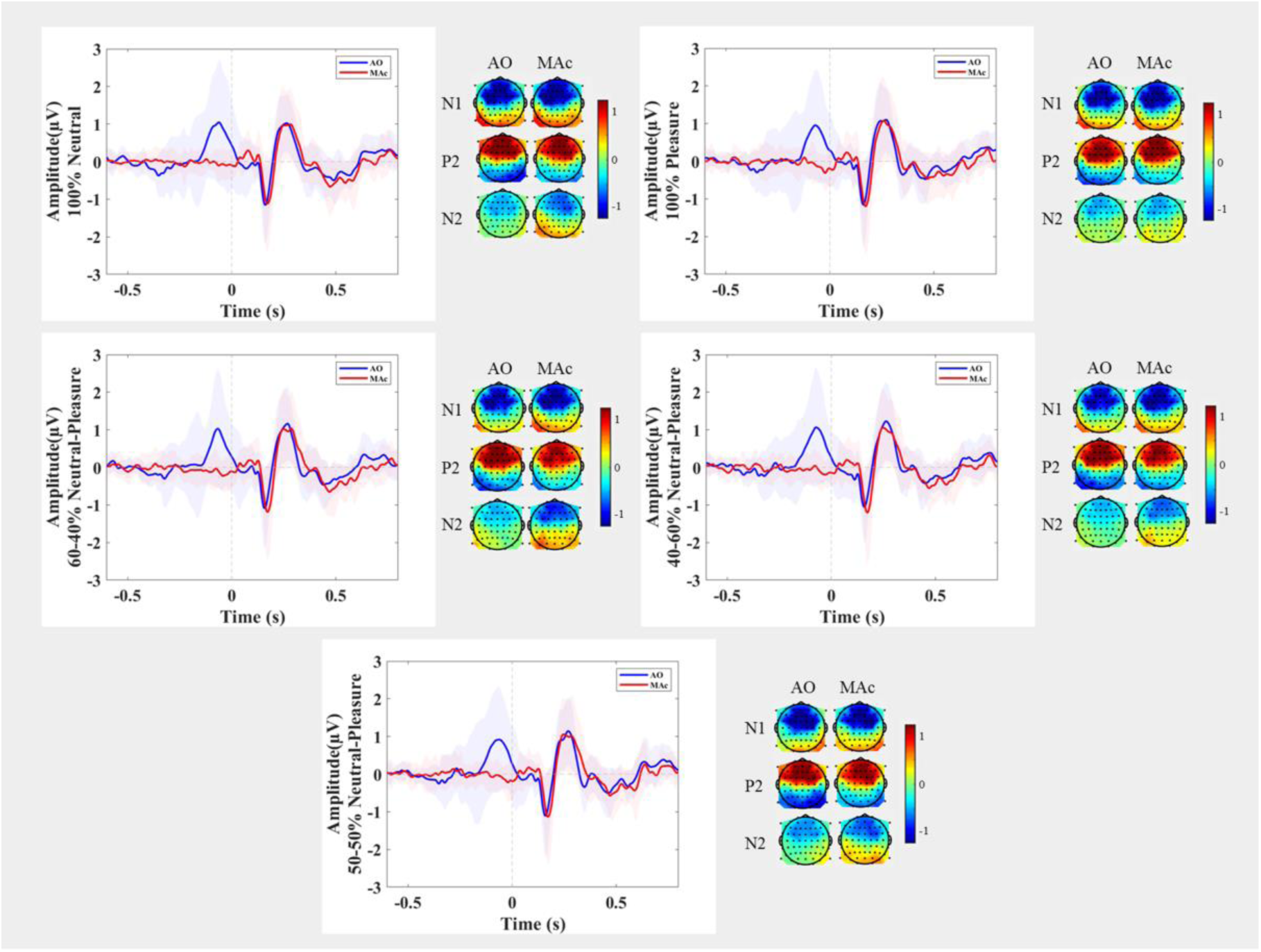
Grand average ERP waveforms ± standard error of mean and topographic maps showing voltage distribution at the peak ERPs, comparing self-generated and externally-generated voices for the five self-voice types originating from a fronto-central ROI. Abbreviations: MAc = Motor Auditory Corrected; AO = Auditory Only.

### Statistical analyses

Statistical analyses on N100, P200 and N200 data were performed in R version 4.2.2 (2022-10-31) Copyright (C) 2022, using linear mixed modeling with lmer and lmerTest packages (Bates, 2016; Kuznetsova, Brockhoff, & Christensen, 2017). We used linear mixed modeling (LMM) to control for the random effects of participants influencing the outcome measure. Additionally, since HP measured by the LSHS is a continuous variable, LMMs are considered more appropriate than classical ANOVA to analyze the impact of HP on sensory prediction (condition) and salience (stimulus type). Amplitude values of the event related potential (N100/P200/N200) were used as an outcome measure, participants were used as random effects, and condition (2 levels: MAc and AO), stimulus type (5 levels: 100% neutral, 60-40% neutral-pleasure, 50-50% neutral-pleasure, 40-60% neutral-pleasure, 100% pleasure) and LSHS total or LSHS AVH scores (continuous variable) were included as fixed effects, respectively, in the hypothesized models. For all the models, the Gaussian distribution of model residuals and quantile-quantile plots confirmed their respective adequacy.

## 3. Results

We followed a hypothesis-driven approach to specifically probe the interaction of sensory feedback processing (condition) and emotional voice quality (stimulus type) with HP.

*N100 & P200:* None of the models testing the interaction of HP with sensory feedback processing (condition) and emotional quality (stimulus type) showed significant differences from the null models (supplementary table 2 and 3).

*N200:* The model that showed best goodness of fit [m1.1_N200 <-lmer(N200 ∼ + Condition * LSHS total + Stimulus Type + (1|ID), data=data, REML = FALSE)] also yielded a significant difference (χ2(7) = 34.621, p = 0.000 **; AIC = 245.43; table 1, figure 3) against the null model [m0_N200; AIC = 266.05]. There was no notable impact of the condition, and the influence of HP on the N200 suppression effect was not statistically significant.

**Figure 3:**
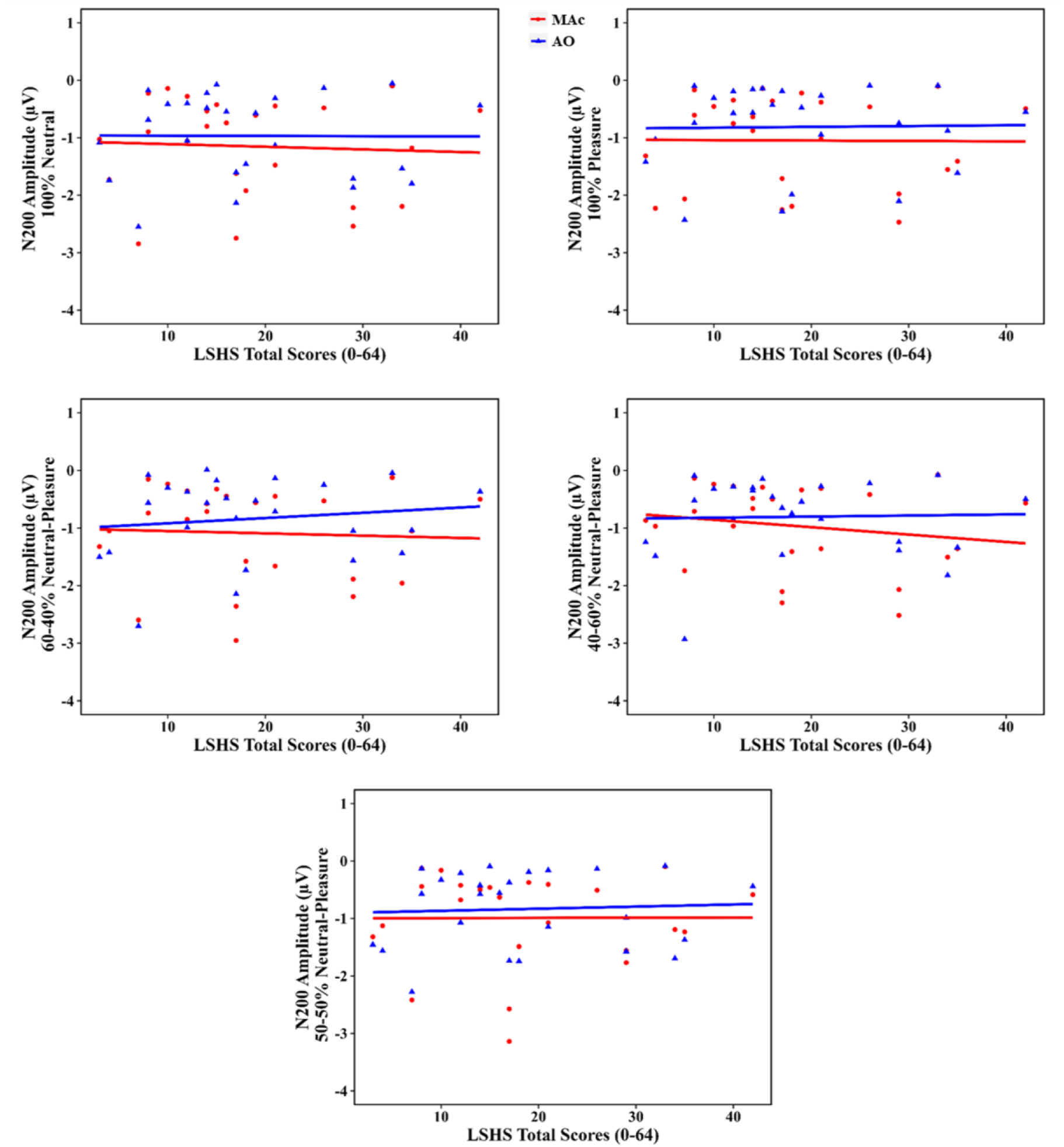
Scatter plots depicting the change in N200 amplitudes as a function of HP (based on LSHS total scores) for each stimulus type. While it appeared as if the N200 response differed for self-and externally-generated self-voices with increase in HP, this result only approached statistical significance (p = 0.056).

**Table 1:** Linear mixed effects model of N200 amplitude including the effect of hallucination proneness based on LSHS total scores. Notes: SE = standard error; SD = standard deviation; *p < 0.05; **p < 0.01; ***p < 0.001. Degrees of freedom for Fixed Effects: df = 225.0 (except Intercept: df = 26.07).

| Variable | Estimate | SE | t value | Pr(>\|t\|) |
| --- | --- | --- | --- | --- |
| **Fixed Effects** |  |  |  |  |
| Intercept | -1.077133 | 0.287934 | -3.741 | 0.000871 *** |
| AO | 0.057689 | 0.082453 | 0.700 | 0.484863 |
| LSHS total | -0.004347 | 0.013474 | -0.323 | 0.749531 |
| 60N | 0.096563 | 0.062670 | 1.541 | 0.124767 |
| 50N | 0.145844 | 0.062670 | 2.327 | 0.020845 * |
| 40N | 0.174131 | 0.062670 | 2.779 | 0.005922 ** |

| Pleasure | 0.128594 | 0.062670 | 2.052 | 0.041335 * |
| --- | --- | --- | --- | --- |
| AO*LSHS total | 0.007476 | 0.003896 | 1.919 | 0.056223 . |
| Groups | Name | Variance | SD |  |
| **Random Effects** |  |  |  |  |
| Subjects | Intercept | 0.45025 | 0.6710 |  |
| Residual |  | 0.09819 | 0.3134 |  |
| Number of observations: 250, Subjects: 25 |  |  |  |  |

## 4. Discussion

This EEG study investigated changes in sensory feedback processing of one’s own voice and attention allocation as a function of HP by manipulating the positive emotionality of self-voice stimuli. Specifically, we examined the auditory N100, P200, and N200 responses for the self-and externally-generated self-voice, using a modified version of a previously established motor-auditory paradigm (figure 1; (Pinheiro et al., 2020; Pinheiro et al., 2018)). However, contrary to expectations, the ERP responses for self-and externally-generated voices were similar, leading to no discernable suppression effects. Moreover, HP did not modulate N100, P200 or N200 responses for the self-and externally-generated voices. Our findings thus question the notion of changes in attentional engagement or sensory feedback processing of the self-voice along the neutral-pleasure emotion expression continuum as a function of HP.

The recognition of vocal emotions relies on identifying physical characteristics such as base frequency, intensity, duration and pitch variability (Banse & Scherer, 1996; Juslin & Laukka, 2001; Kanske, Schonfelder, & Wessa, 2013; Sauter, Eisner, Calder, & Scott, 2010). Specifically, vocal emotions characterized by moderate or low emotional intensity lead to diminished accuracy in recognizing emotions and require a longer processing time (Banse & Scherer, 1996; Juslin & Laukka, 2001; Kanske et al., 2013; Sauter, 2017; Sauter et al., 2010). For example, pleasure vocalizations are most often associated with long duration, low spectral center of gravity and high spectral variation (Sauter et al., 2010). These characteristics may pose challenges for capturing the impact of these vocalizations in early ERPs. Further, pleasure vocalizations are often confused with other positive emotions such as relief and contentment, leading to unclear and ambiguous overall perceptions (Sauter, 2017; Sauter et al., 2010). Notably, these emotions share similar physical properties as well as arousal and valence ratings, even though they differ semantically, meaning they are not synonyms (Sauter, 2017; Sauter et al., 2010). Based on these aspects, it is likely that the perceptual discriminability among the five types of self-vocalizations varying in pleasure content used here might have been low. Further, using the same sample of participants and design but a self-voice continuum from neutral to angry yielded a significant global N100 suppression effect (Duggirala et al., 2023). Hence, the question arises as to why similar effects did not emerge as a function of a continuum manipulation for positive vocal emotion expressions. Participants in the current study did not exhibit a complete sense of ownership (“my voice or someone else’s voice”) and agency (“feeling associated with the sensory outcome of one’s voluntary action”) of their own 100% pleasure voice. The sense of ownership was lower for the 100% pleasure stimuli compared to the 100% angry self-voice ((Herbert, Herbert, Ethofer, & Pauli, 2011; Yoshie & Haggard, 2013); supplementary figure 1C and supplementary figure 1C of (Duggirala et al., 2023)). Likewise, in the epistemological domain of positive emotions, pleasure vocalizations link to a comparatively diminished sense of agency when contrasted with elation or pride (Sauter, 2017). Likewise, research indicates that emotion perception is susceptible to contextual influence (Liuni, Ponsot, Bryant, & Aucouturier, 2020; Mauchand & Zhang, 2023; Paulmann & Pell, 2010). It is likely that the prevalence of heightened ambiguous expression (3:2 → ambiguous:unambiguous self-voice), next to perceptual discriminability, might have created an overall ambiguous context. Therefore, the lack of perceptual discriminability among the self-voices from the neutral-pleasure continuum and the low sense of ownership and agency, might have established an overall ambiguous context. In turn, these factors might have resulted in no changes in the certainty of sensory feedback in self-voice or differential attentional engagement, leading to no global N100/P200 suppression effect and no discernable effects of voice quality on N100/P200 suppression effects.

Research findings indicate that the processing and attention-capturing functions of positive emotions require a longer duration and are evidenced in the ERP components occurring in approximately the latency range of 200-550 ms (Carretié, Hinojosa, Martín-Loeches, Mercado, & Tapia, 2004; Carretié, Martín-Loeches, Hinojosa, & Mercado, 2001; Xue et al., 2013). The N200 response (regardless of condition and HP; table 1, supplementary table 2, supplementary table 5) showed a global reduction for self-voices with more than 50% pleasure content. Further, while it appeared as if the N200 response differed for self-and externally-generated self-voices with increasing HP, this result only approached statistical significance (p = 0.056, table 1, figure 3). Considering the initial argument of lack of ownership in pleasure vocalizations and ambiguous context, this reduced N200 response for self-voices high on pleasure content, might reflect a decreased ability to categorize and process these voices in general (compared to 100% neutral voice) but decreased further in high HP individuals. Alternatively, this might also indicate an absence of attentional bias towards pleasure compared to non-emotional neutral expressions. Previous studies have shown that schizotypy is associated with less attentional focus on positive affect and feelings, and experiencing and anticipating less pleasure (Giakoumaki, 2016; Kerns, Docherty, & Martin, 2008; Li et al., 2019; Martin, Becker, Cicero, Docherty, & Kerns, 2011). Likewise, individuals scoring high on HP misidentified pleasure content in vocalizations from the pleasure-neutral continuum (Amorim et al., 2022). Authors in this study have speculated that the high HP participants’ intrinsic attentional bias towards negative emotions leading them to possibly ignore or allocate less attentional resources to non-negative emotional stimuli, i.e., neutral or pleasure.

A limitation related to the task-design is noted. In contrast to the conventional blocked design in auditory-motor tasks, the current mixed design included a visual cue to prompt the respective condition of interest. Furthermore, there was a fixed duration of 500 ms between the visual cue and the onset of the auditory stimulus in the externally-generated condition. While the influence of the visual cue was eliminated in the MA condition by subtracting MO from MA to eliminate motor effects, in the AO condition, the response to the visual cue remained as a pre-stimulus positive potential. Both the fixed delay and the visual cue could have enhanced anticipation of the externally-generated voice and focused attention on it (Heynckes, De Weerd, Valente, Formisano, & De Martino, 2020; Sowman, Kuusik, & Johnson, 2012). Consequently, the listener might have been better prepared for the sound to occur, approximating the preparedness for the sensory consequences of a self-generated voice in the MA condition (Costa-Faidella, Baldeweg, Grimm, & Escera, 2011; Heynckes et al., 2020; Sowman et al., 2012). However, it should be noted that previous studies using both cued and uncued externally-generated conditions showed no difference in suppression effects (Griffiths et al., 2023; Lange, 2013). Studies have reported that it is not the motor-action per se, but the voluntary intention involving motor planning to self-generate an action (e.g., a voice) that leads to a sensory suppression (Jack et al., 2021; Timm, SanMiguel, Keil, Schroger, & Schonwiesner, 2014). Therefore, the null findings associated with the N100/P200 suppression effects cannot be attributed exclusively to factors associated with the current task-design.

Various factors (e.g., lack of acoustic differences, ambiguous context, task design) discussed earlier are probable contributors to the absence of significant findings in the present study. Future investigations should take these factors into account and analyze voices in distinct blocks. Despite the lack of significant results in this study, the investigation of positive emotions in processing a self-relevant stimulus (e.g., self-voice) is important. Voice hearers with a psychotic disorder have difficulties in experiencing positive emotions such as pleasure and motivation (Cohen & Minor, 2010; Horan et al., 2008; Kring & Moran, 2008; Li et al., 2019; Watson & Naragon-Gainey, 2010). These difficulties are associated with negative symptoms such as constricted affect and social aloofness, which in turn are influenced by positive symptoms such as hallucinations (Watson & Naragon-Gainey, 2010). Therefore, probing how processing of a positive self-relevant stimulus (e.g., self-voice) affect allocation of attention and sensory feedback processing of own voice in non-clinical individuals who are highly prone to hallucination can contribute to understanding the pre-existing limitations in processing positive emotions within non-clinical samples. Future studies should also include assessments of negative symptoms (e.g., lack of motivation, depression, stress) while studying positive emotions in voice hearing as they may act as a mediating factor in the relationship between positive emotion processing and HP. This, in turn, may provide insights for developing approaches to enhance pleasure perception and increase motivation in voice hearers.

## Data availability

The data that support the findings of this study are available from the corresponding author upon reasonable request.

## Author Contributions

SXD, MS, DL, AP, SK conceptualized and designed the experiment, SXD prepared materials, collected and analyzed the data, and wrote the first draft of the manuscript, SXD, MS, DL, AP, SK refined the manuscript. MS, AP, SK procured funding for the project. All authors have approved the final version of the manuscript.

## Acknowledgements

The authors would like to thank Joseph Johnson for sharing the paradigm code that was adapted and used for the current experiment. We would also like to thank Alexandra Emmendorfer and Radhika Rajan for detailed discussions on data analysis and trouble shooting matlab code.

## References

Addington, J., Penn, D., Woods, S. W., Addington, D., & Perkins, D. O. (2008). Facial affect recognition in individuals at clinical high risk for psychosis. Br J Psychiatry, 192(1), 67–68. doi:10.1192/bjp.bp.107.039784

Alba-Ferrara, L., de Erausquin, G. A., Hirnstein, M., Weis, S., & Hausmann, M. (2013). Emotional prosody modulates attention in schizophrenia patients with hallucinations. Front Hum Neurosci, 7, 59. doi:10.3389/fnhum.2013.00059

Allen, P., Aleman, A., & McGuire, P. K. (2007). Inner speech models of auditory verbal hallucinations: evidence from behavioural and neuroimaging studies. Int Rev Psychiatry, 19(4), 407–415. doi: 10.1080/09540260701486498

Amminger, G. P., Schafer, M. R., Klier, C. M., Schlogelhofer, M., Mossaheb, N., Thompson, A., . . . Nelson, B. (2012). Facial and vocal affect perception in people at ultra-high risk of psychosis, first-episode schizophrenia and healthy controls. Early Interv Psychiatry, 6(4), 450–454. doi: 10.1111/j.1751-7893.2012.00362.x

Amminger, G. P., Schafer, M. R., Papageorgiou, K., Klier, C. M., Schlogelhofer, M., Mossaheb, N., . . . McGorry, P. D. (2012). Emotion recognition in individuals at clinical high-risk for schizophrenia. Schizophr Bull, 38(5), 1030–1039. doi: 10.1093/schbul/sbr015

Amorim, M., Roberto, M. S., Kotz, S. A., & Pinheiro, A. P. (2022). The perceived salience of vocal emotions is dampened in non-clinical auditory verbal hallucinations. Cogn Neuropsychiatry, 27(2-3), 169–182. doi: 10.1080/13546805.2021.1949972

Baess, P., Widmann, A., Roye, A., Schroger, E., & Jacobsen, T. (2009). Attenuated human auditory middle latency response and evoked 40-Hz response to self-initiated sounds. Eur J Neurosci, 29(7), 1514–1521. doi: 10.1111/j.1460-9568.2009.06683.x

Banse, R., & Scherer, K. R. (1996). Acoustic profiles in vocal emotion expression. J Pers Soc Psychol, 70(3), 614.

Barkus, E., Stirling, J., Hopkins, R., McKie, S., & Lewis, S. (2007). Cognitive and neural processes in non-clinical auditory hallucinations. Br J Psychiatry Suppl, 51, s76–81. doi: 10.1192/bjp.191.51.s76

Bates, D. (2016). lme4: Linear mixed-effects models using Eigen and S4. R package version, 1, 1.

Baumeister, D., Sedgwick, O., Howes, O., & Peters, E. (2017). Auditory verbal hallucinations and continuum models of psychosis: A systematic review of the healthy voice-hearer literature. Clin Psychol Rev, 51, 125–141. doi: 10.1016/j.cpr.2016.10.010

Behroozmand, R., Karvelis, L., Liu, H., & Larson, C. R. (2009). Vocalization-induced enhancement of the auditory cortex responsiveness during voice F0 feedback perturbation. Clin Neurophysiol, 120(7), 1303–1312. doi: 10.1016/j.clinph.2009.04.022

Birchwood, M., & Chadwick, P. (1997). The omnipotence of voices: testing the validity of a cognitive model. Psychol Med, 27(6), 1345–1353.

Blakemore, S. J., Wolpert, D., & Frith, C. (2000). Why can’t you tickle yourself? Neuroreport, 11(11), R11–16. doi: 10.1097/00001756-200008030-00002

Brebion, G., Stephan-Otto, C., Ochoa, S., Roca, M., Nieto, L., & Usall, J. (2016). Impaired Self-Monitoring of Inner Speech in Schizophrenia Patients with Verbal Hallucinations and in Non-clinical Individuals Prone to Hallucinations. Front Psychol, 7, 1381. doi: 10.3389/fpsyg.2016.01381

Carretié, L., Hinojosa, J. A., Martín-Loeches, M., Mercado, F., & Tapia, M. (2004). Automatic attention to emotional stimuli: neural correlates. Hum Brain Mapp, 22(4), 290–299.

Carretié, L., Martín-Loeches, M., Hinojosa, J. A., & Mercado, F. (2001). Emotion and attention interaction studied through event-related potentials. J Cogn Neurosci, 13(8), 1109–1128.

Castiajo, P., & Pinheiro, A. P. (2017). On “Hearing” Voices and “Seeing” Things: Probing Hallucination Predisposition in a Portuguese Nonclinical Sample with the Launay-Slade Hallucination Scale-Revised. Front Psychol, 8, 1138. doi: 10.3389/fpsyg.2017.01138

Chen, Z., Chen, X., Liu, P., Huang, D., & Liu, H. (2012). Effect of temporal predictability on the neural processing of self-triggered auditory stimulation during vocalization. BMC Neurosci, 13, 55. doi: 10.1186/1471-2202-13-55

Cohen, A. S., & Minor, K. S. (2010). Emotional experience in patients with schizophrenia revisited: meta-analysis of laboratory studies. Schizophrenia Bulletin, 36(1), 143–150.

Cook, S., & Wilding, J. (2001). Earwitness testimony: effects of exposure and attention on the face overshadowing effect. Br J Psychol, 92(Pt 4), 617–629. doi: 10.1348/000712601162374

Costa-Faidella, J., Baldeweg, T., Grimm, S., & Escera, C. (2011). Interactions between “what” and “when” in the auditory system: temporal predictability enhances repetition suppression. Journal of Neuroscience, 31(50), 18590–18597.

Daalman, K., Boks, M. P., Diederen, K. M., de Weijer, A. D., Blom, J. D., Kahn, R. S., & Sommer, I. E. (2011). The same or different? A phenomenological comparison of auditory verbal hallucinations in healthy and psychotic individuals. J Clin Psychiatry, 72(3), 320–325. doi: 10.4088/JCP.09m05797yel

Diederen, K. M., Daalman, K., de Weijer, A. D., Neggers, S. F., van Gastel, W., Blom, J. D., . . . Sommer, I. E. (2012). Auditory hallucinations elicit similar brain activation in psychotic and nonpsychotic individuals. Schizophr Bull, 38(5), 1074–1082. doi: 10.1093/schbul/sbr033

Duggirala, S. X., Schwartze, M., Goller, L. K., Linden, D. E., Pinheiro, A., & Kotz, S. A. (2023). Hallucination proneness alters sensory feedback processing in self-voice production. bioRxiv, 2023.2007. 2028.550971.

Feinberg, I. (1978). Efference copy and corollary discharge: implications for thinking and its disorders. Schizophrenia Bulletin, 4(4), 636.

Ford, J. M., & Mathalon, D. H. (2004). Electrophysiological evidence of corollary discharge dysfunction in schizophrenia during talking and thinking. J Psychiatr Res, 38(1), 37–46. doi: 10.1016/s0022-3956(03)00095-5

Ford, J. M., Mathalon, D. H., Heinks, T., Kalba, S., Faustman, W. O., & Roth, W. T. (2001). Neurophysiological evidence of corollary discharge dysfunction in schizophrenia. Am J Psychiatry, 158(12), 2069–2071. doi: 10.1176/appi.ajp.158.12.2069

Ford, J. M., Mathalon, D. H., Kalba, S., Whitfield, S., Faustman, W. O., & Roth, W. T. (2001a). Cortical responsiveness during inner speech in schizophrenia: an event-related potential study. Am J Psychiatry, 158(11), 1914–1916. doi: 10.1176/appi.ajp.158.11.1914

Ford, J. M., Mathalon, D. H., Kalba, S., Whitfield, S., Faustman, W. O., & Roth, W. T. (2001b). Cortical responsiveness during talking and listening in schizophrenia: an event-related brain potential study. Biol Psychiatry, 50(7), 540–549. doi: 10.1016/s0006-3223(01)01166-0

Ford, J. M., Mathalon, D. H., Roach, B. J., Keedy, S. K., Reilly, J. L., Gershon, E. S., & Sweeney, J. A. (2013). Neurophysiological evidence of corollary discharge function during vocalization in psychotic patients and their nonpsychotic first-degree relatives. Schizophr Bull, 39(6), 1272–1280. doi: 10.1093/schbul/sbs129

Foussias, G., Agid, O., Fervaha, G., & Remington, G. (2014). Negative symptoms of schizophrenia: clinical features, relevance to real world functioning and specificity versus other CNS disorders. European Neuropsychopharmacology, 24(5), 693–709.

Frith, C., Friston, K., Liddle, P., & Frackowiak, R. (1992). PET imaging and cognition in schizophrenia. Journal of the Royal Society of Medicine, 85(4), 222–224.

Frith, C. D., Blakemore, S., & Wolpert, D. M. (2000). Explaining the symptoms of schizophrenia: abnormalities in the awareness of action. Brain Res Brain Res Rev, 31(2-3), 357–363. doi: 10.1016/s0165-0173(99)00052-1

Galdos, M., Simons, C., Fernandez-Rivas, A., Wichers, M., Peralta, C., Lataster, T., . . . van Os, J. (2011). Affectively salient meaning in random noise: a task sensitive to psychosis liability. Schizophr Bull, 37(6), 1179–1186. doi: 10.1093/schbul/sbq029

Giakoumaki, S. G. (2016). Emotion processing deficits in the different dimensions of psychometric schizotypy. Scand J Psychol, 57(3), 256–270.

Griffiths, O., Jack, B. N., Pearson, D., Elijah, R., Mifsud, N., Han, N., . . . Whitford, T. J. (2023). Disrupted auditory N1, theta power and coherence suppression to willed speech in people with schizophrenia. Neuroimage Clin, 37, 103290. doi: 10.1016/j.nicl.2022.103290

Herbert, C., Herbert, B. M., Ethofer, T., & Pauli, P. (2011). His or mine? The time course of self– other discrimination in emotion processing. Soc Neurosci, 6(3), 277–288.

Heynckes, M., De Weerd, P., Valente, G., Formisano, E., & De Martino, F. (2020). Behavioral effects of rhythm, carrier frequency and temporal cueing on the perception of sound sequences. PLoS One, 15(6), e0234251.

Horan, W. P., Blanchard, J. J., Clark, L. A., & Green, M. F. (2008). Affective traits in schizophrenia and schizotypy. Schizophrenia Bulletin, 34(5), 856–874.

Hughes, G., Desantis, A., & Waszak, F. (2013). Mechanisms of intentional binding and sensory attenuation: the role of temporal prediction, temporal control, identity prediction, and motor prediction. Psychol Bull, 139(1), 133–151. doi: 10.1037/a0028566

Jack, B. N., Chilver, M. R., Vickery, R. M., Birznieks, I., Krstanoska-Blazeska, K., Whitford, T. J., & Griffiths, O. (2021). Movement planning determines sensory suppression: An event-related potential study. J Cogn Neurosci, 33(12), 2427–2439.

Johns, L. C., Hemsley, D., & Kuipers, E. (2002). A comparison of auditory hallucinations in a psychiatric and non-psychiatric group. Br J Clin Psychol, 41(Pt 1), 81–86.

Johns, L. C., Kompus, K., Connell, M., Humpston, C., Lincoln, T. M., Longden, E., . . . Laroi, F. (2014). Auditory verbal hallucinations in persons with and without a need for care. Schizophr Bull, 40 Suppl 4(Suppl 4), S255-264. doi: 10.1093/schbul/sbu005

Johnson, M. K., Hashtroudi, S., & Lindsay, D. S. (1993). Source monitoring. Psychol Bull, 114(1), 3–28. doi: 10.1037/0033-2909.114.1.3

Jones, S. R., & Fernyhough, C. (2007). Thought as action: Inner speech, self-monitoring, and auditory verbal hallucinations. Conscious Cogn, 16(2), 391–399.

Juslin, P. N., & Laukka, P. (2001). Impact of intended emotion intensity on cue utilization and decoding accuracy in vocal expression of emotion. Emotion, 1(4), 381.

Kanske, P., Schonfelder, S., & Wessa, M. (2013). Emotional modulation of the attentional blink and the relation to interpersonal reactivity. Front Hum Neurosci, 7, 641. doi: 10.3389/fnhum.2013.00641

Kawahara, H. (2006). STRAIGHT, exploitation of the other aspect of VOCODER: Perceptually isomorphic decomposition of speech sounds. Acoustical science and technology, 27(6), 349–353.

Kawahara, H., & Irino, T. (2005). Underlying principles of a high-quality speech manipulation system STRAIGHT and its application to speech segregation Speech separation by humans and machines (pp. 167–180): Springer.

Kawahara, H., Morise, M., Takahashi, T., Nisimura, R., Irino, T., & Banno, H. (2008). Tandem-STRAIGHT: A temporally stable power spectral representation for periodic signals and applications to interference-free spectrum, F0, and aperiodicity estimation. Paper presented at the 2008 IEEE International Conference on Acoustics, Speech and Signal Processing.

Kerns, J. G., Docherty, A. R., & Martin, E. A. (2008). Social and physical anhedonia and valence and arousal aspects of emotional experience. J Abnorm Psychol, 117(4), 735.

Knolle, F., Schwartze, M., Schroger, E., & Kotz, S. A. (2019). Auditory Predictions and Prediction Errors in Response to Self-Initiated Vowels. Front Neurosci, 13, 1146. doi: 10.3389/fnins.2019.01146

Korzyukov, O., Karvelis, L., Behroozmand, R., & Larson, C. R. (2012). ERP correlates of auditory processing during automatic correction of unexpected perturbations in voice auditory feedback. Int J Psychophysiol, 83(1), 71–78. doi: 10.1016/j.ijpsycho.2011.10.006

Kring, A. M., & Moran, E. K. (2008). Emotional response deficits in schizophrenia: insights from affective science. Schizophrenia Bulletin, 34(5), 819–834.

Kuznetsova, A., Brockhoff, P. B., & Christensen, R. H. (2017). lmerTest package: tests in linear mixed effects models. Journal of statistical software, 82, 1–26.

Lange, K. (2013). The ups and downs of temporal orienting: a review of auditory temporal orienting studies and a model associating the heterogeneous findings on the auditory N1 with opposite effects of attention and prediction. Front Hum Neurosci, 7, 263. doi: 10.3389/fnhum.2013.00263

Larøi, F. (2012). How do auditory verbal hallucinations in patients differ from those in non-patients? Front Hum Neurosci, 6, 25.

Larøi, F., & Van der Linden, M. (2005a). Metacognitions in proneness towards hallucinations and delusions. Behaviour research and Therapy, 43(11), 1425–1441.

Larøi, F., & Van Der Linden, M. (2005b). Nonclinical Participants’ Reports of Hallucinatory Experiences. Canadian Journal of Behavioural Science/Revue canadienne des sciences du comportement, 37(1), 33.

Launay, G., & Slade, P. (1981). The measurement of hallucinatory predisposition in male and female prisoners. Personality and Individual Differences, 2(3), 221–234.

Li, L. Y., Fung, C. K., Moore, M. M., & Martin, E. A. (2019). Differential emotional abnormalities among schizotypy clusters. Schizophr Res. doi: 10.1016/j.schres.2019.01.042

Liuni, M., Ponsot, E., Bryant, G. A., & Aucouturier, J.-J. (2020). Sound context modulates perceived vocal emotion. Behavioural processes, 172, 104042.

Maijer, K., Begemann, M. J. H., Palmen, S., Leucht, S., & Sommer, I. E. C. (2018). Auditory hallucinations across the lifespan: a systematic review and meta-analysis. Psychol Med, 48(6), 879–888. doi: 10.1017/S0033291717002367

Martin, E. A., Becker, T. M., Cicero, D. C., Docherty, A. R., & Kerns, J. G. (2011). Differential associations between schizotypy facets and emotion traits. Psychiatry Res, 187(1-2), 94–99. doi: 10.1016/j.psychres.2010.12.028

Mauchand, M., & Zhang, S. (2023). Disentangling emotional signals in the brain: an ALE meta-analysis of vocal affect perception. *Cognitive, Affective*, & Behavioral Neuroscience, 23(1), 17–29.

Mohanty, A., Heller, W., Koven, N. S., Fisher, J. E., Herrington, J. D., & Miller, G. A. (2008). Specificity of emotion-related effects on attentional processing in schizotypy. Schizophr Res, 103(1-3), 129–137. doi: 10.1016/j.schres.2008.03.003

Nelson, B., Whitford, T. J., Lavoie, S., & Sass, L. A. (2014a). What are the neurocognitive correlates of basic self-disturbance in schizophrenia? Integrating phenomenology and neurocognition: Part 2 (aberrant salience). Schizophr Res, 152(1), 20–27. doi: 10.1016/j.schres.2013.06.033

Nelson, B., Whitford, T. J., Lavoie, S., & Sass, L. A. (2014b). What are the neurocognitive correlates of basic self-disturbance in schizophrenia?: Integrating phenomenology and neurocognition. Part 1 (Source monitoring deficits). Schizophr Res, 152(1), 12–19. doi: 10.1016/j.schres.2013.06.022

Nguyen, A., Frobert, L., McCluskey, I., Golay, P., Bonsack, C., & Favrod, J. (2016). Development of the positive emotions program for schizophrenia: an intervention to improve pleasure and motivation in schizophrenia. Front Psychiatry, 7, 13.

Paulmann, S., & Pell, M. D. (2010). Dynamic emotion processing in Parkinson’s disease as a function of channel availability. J Clin Exp Neuropsychol, 32(8), 822–835. doi: 10.1080/13803391003596371

Pinheiro, A. P., Schwartze, M., Amorim, M., Coentre, R., Levy, P., & Kotz, S. A. (2020). Changes in motor preparation affect the sensory consequences of voice production in voice hearers. Neuropsychologia, 146, 107531. doi: 10.1016/j.neuropsychologia.2020.107531

Pinheiro, A. P., Schwartze, M., & Kotz, S. A. (2018). Voice-selective prediction alterations in nonclinical voice hearers. Sci Rep, 8(1), 14717. doi: 10.1038/s41598-018-32614-9

Rossell, S. L., & Boundy, C. L. (2005). Are auditory-verbal hallucinations associated with auditory affective processing deficits? Schizophr Res, 78(1), 95–106. doi: 10.1016/j.schres.2005.06.002

Sauter, D. A. (2017). The nonverbal communication of positive emotions: An emotion family approach. Emotion Review, 9(3), 222–234.

Sauter, D. A., Eisner, F., Calder, A. J., & Scott, S. K. (2010). Perceptual cues in nonverbal vocal expressions of emotion. Q J Exp Psychol (Hove), 63(11), 2251–2272.

Schweinberger, S. R., Herholz, A., & Sommer, W. (1997). Recognizing famous voices: influence of stimulus duration and different types of retrieval cues. J Speech Lang Hear Res, 40(2), 453–463. doi: 10.1044/jslhr.4002.453

Sowman, P. F., Kuusik, A., & Johnson, B. W. (2012). Self-initiation and temporal cueing of monaural tones reduce the auditory N1 and P2. Exp Brain Res, 222, 149–157.

Swink, S., & Stuart, A. (2012a). Auditory long latency responses to tonal and speech stimuli. J Speech Lang Hear Res, 55(2), 447–459. doi: 10.1044/1092-4388(2011/10-0364)

Swink, S., & Stuart, A. (2012b). The effect of gender on the N1-P2 auditory complex while listening and speaking with altered auditory feedback. Brain Lang, 122(1), 25–33. doi: 10.1016/j.bandl.2012.04.007

Timm, J., SanMiguel, I., Keil, J., Schroger, E., & Schonwiesner, M. (2014). Motor intention determines sensory attenuation of brain responses to self-initiated sounds. J Cogn Neurosci, 26(7), 1481–1489. doi: 10.1162/jocn_a_00552

van ‘t Wout, M., Aleman, A., Kessels, R. P., Laroi, F., & Kahn, R. S. (2004). Emotional processing in a non-clinical psychosis-prone sample. Schizophr Res, 68(2-3), 271–281. doi: 10.1016/j.schres.2003.09.006

van Os, J. (2003). Is there a continuum of psychotic experiences in the general population? Epidemiol Psichiatr Soc, 12(4), 242–252. doi: 10.1017/s1121189x00003067

van Os, J., Linscott, R. J., Myin-Germeys, I., Delespaul, P., & Krabbendam, L. (2009). A systematic review and meta-analysis of the psychosis continuum: evidence for a psychosis proneness-persistence-impairment model of psychotic disorder. Psychol Med, 39(2), 179–195. doi: 10.1017/S0033291708003814

Ventura, M. I., Nagarajan, S. S., & Houde, J. F. (2009). Speech target modulates speaking induced suppression in auditory cortex. BMC Neurosci, 10(1), 1–11.

Watson, D., & Naragon-Gainey, K. (2010). On the specificity of positive emotional dysfunction in psychopathology: Evidence from the mood and anxiety disorders and schizophrenia/schizotypy. Clin Psychol Rev, 30(7), 839–848.

Wolpert, D. M., Ghahramani, Z., & Jordan, M. I. (1995). An internal model for sensorimotor integration. Science, 269(5232), 1880–1882. doi: 10.1126/science.7569931

Xue, S., Cui, J., Wang, K., Zhang, S., Qiu, J., & Luo, Y. (2013). Positive emotion modulates cognitive control: an event-related potentials study. Scand J Psychol, 54(2), 82–88.

Yoshie, M., & Haggard, P. (2013). Negative emotional outcomes attenuate sense of agency over voluntary actions. Current Biology, 23(20), 2028–2032.

