## supplementary document for "Not enough pleasure? Influence of hallucination proneness on sensory feedback processing of positive self-voice"

**Table and table legends**

**Supplementary table 1:** Neutral-Pleasure continua with 11 voice morphs.

a) Neutral-to-pleasure

| Emotion / Morphs | **1*** | 2 | 3 | 4 | **5*** | **6*** | 7* | 8 | 9 | 10 | **11*** |
|---|---|---|---|---|---|---|---|---|---|---|---|
| Neutral | **100 %** | 90 % | 80 % | 70 % | **60 %** | **50 %** | 40 % | 30 % | 20 % | 10 % | **0%** |
| Pleasure | **0%** | 10 % | 20 % | 30 % | **40 %** | **50 %** | 60 % | 70 % | 80 % | 90 % | **100 %** |

b) Angry-to-pleasure

| Emotion / Morphs | **1*** | 2 | 3 | 4 | **5*** | **6*** | 7* | 8 | 9 | 10 | **11*** |
|---|---|---|---|---|---|---|---|---|---|---|---|
| Pleasure | **100 %** | 90 % | 80 % | 70 % | **60 %** | **50 %** | 40 % | 30 % | 20 % | 10 % | **0%** |
| Neutral | **0%** | 10 % | 20 % | 30 % | **40 %** | **50 %** | 60 % | 70 % | 80 % | 90 % | **100 %** |

c) Final Stimuli for Ah and Oh vocalizations.

| |
|---|
| 100% Neutral = Ah (a1 + b11) + Oh (a1 + b11) |
| 60-40% Neutral-Pleasure = Ah (a5 + b7) + Oh (a5 + b7) |
| 50-50% Neutral-Pleasure= Ah (a6 + b6) + Oh (a6 + b6) |

| |
|---|
| 40-60% Neutral-Pleasure = Ah (a7 + b5) + Oh (a7 + b5) |
| 100% Pleasure = Ah (a11 + b1) + Oh (a11 + b1) |

Note: a1 refers to the specific voice morph from table a, voice morph 1.

**Supplementary table 2:** Mean latencies of N100, P200 and N200 amplitudes.

| | Mean (ms) | S.d (ms) | Min (ms) | Max (ms) |
|---|---|---|---|---|
| N100 | 162 | 0.027 | 0.08 | 0.23 |
| P200 | 265 | 0.033 | 0.18 | 0.38 |
| N200 | 462 | 0.078 | 0.25 | 0.6 |

**Supplementary table 3:** Model comparisons with N100 amplitude as output and HP, Condition and Stimulus.

Notes: SE = standard error; SD = standard deviation; NP = number of parameters; AIC = Akaike information criterion; BIC = Bayesian information criterion; Chisq = chi square; Df = degree of freedom; M1 = Condition * Stimulus type * LSHS total/AVH; M2 = Condition * LSHS total/AVH + Stimulus type * LSHS total/AVH; M3 = Condition * LSHS total + Stimulus type; *p < 0.05; **p < 0.01; ***p < 0.001.

| **Models with LSHS total** | | | | | | | | |
|---|---|---|---|---|---|---|---|---|
| **Model** | **Parameters** | **AIC** | **BIC** | **Log Lik** | **Deviance** | **Chisq** | **Df** | **Pr(>Chisq)** |
| Null model | 3 | 372.29 | 382.85 | -183.14 | 366.29 | | | |
| M1 | 22 | 402.08 | 479.55 | -179.04 | 358.08 | 8.2081 | 19 | 0.9845 |
| M2 | 14 | 387.77 | 437.07 | -179.89 | 359.77 | 6.5166 | 11 | 0.8368 |
| M3 | 10 | 380.61 | 415.83 | -180.31 | 360.61 | 5.673 | 7 | 0.5784 |

| Models with LSHS AVH | | | | | | | | |
|---|---|---|---|---|---|---|---|---|
| **Model** | **Paramet ers** | **AIC** | **BIC** | **Log Lik** | **Deviance** | **Chisq** | **Df** | **Pr(>Chisq)** |
| M1 | 22 | 400.02 | 477.49 | -178.01 | 356.02 | 10.268 | 19 | 0.946 |
| M2 | 14 | 385.98 | 435.28 | -178.99 | 357.98 | 8.3045 | 11 | 0.6858 |
| M3 | 10 | 378.66 | 413.88 | -179.33 | 358.66 | 7.6243 | 7 | 0.3669 |

**Supplementary table 4:** Model comparisons with P200 amplitude as output and HP, Condition and Stimulus.

Notes: SE = standard error; SD = standard deviation; NP = number of parameters; AIC = Akaike information criterion; BIC = Bayesian information criterion; Chisq = chi square; Df = degree of freedom; M1 = Condition * Stimulus type * LSHS total/AVH; M2 = Condition * LSHS total/AVH + Stimulus type * LSHS total/AVH; M3 = Condition * LSHS total + Stimulus type; *p < 0.05; **p < 0.01; ***p < 0.001.

| Models with LSHS total | | | | | | | | |
|---|---|---|---|---|---|---|---|---|
| **Model** | **Parameters** | **AIC** | **BIC** | **Log Lik** | **Deviance** | **Chisq** | **Df** | **Pr(>Chisq)** |
| Null model | 3 | 332.37 | 342.94 | -163.19 | 326.37 | | | |
| M1 | 22 | 355.08 | 432.56 | -155.54 | 311.08 | 15.288 | 19 | 0.7041 |
| M2 | 14 | 345.81 | 395.11 | -158.91 | 317.81 | 8.5623 | 11 | 0.6622 |
| M3 | 10 | 341.81 | 377.02 | -160.90 | 321.81 | 4.5642 | 7 | 0.713 |

| Models with LSHS AVH | | | | | | | | |
|---|---|---|---|---|---|---|---|---|
| **Model** | **Parameters** | **AIC** | **BIC** | **Log Lik** | **Deviance** | **Chisq** | **Df** | **Pr(>Chisq)** |
| M1 | 22 | 360.75 | 438.23 | -158.38 | 316.75 | 9.6177 | 19 | 0.9618 |
| M2 | 14 | 348.73 | 398.03 | -160.36 | 320.73 | 5.6463 | 11 | 0.8959 |
| M3 | 10 | 341.82 | 377.04 | -160.91 | 321.82 | 4.5476 | 7 | 0.715 |

The influence of hallucination proneness based on LSHS AVH scores was also tested. The model that showed the best goodness of fit [m2.1_N200 <- lmer (N200 ~ + Condition * LSHS AVH + Stimulus Type + (1|ID), data=data, REML = FALSE)] also yielded a significant difference ($\chi2(7) = 31.439$, p = 0.000 **; AIC = 248.61; supplementary table 5, supplementary figure 7) against the null model [m0_N200; AIC = 266.05]. The N200 response for self- compared to externally-generated voices is significantly different, regardless of voice quality or HP.

**Supplementary table 5:** Linear mixed effects model of N200 amplitude including the effect of hallucination proneness based on LSHS AVH scores. Notes: SE = standard error; SD = standard deviation; *p < 0.05; **p < 0.01; ***p < 0.001. Degrees of freedom for Fixed Effects: df = 225.0 (except Intercept: df = 28.54). There was a significant difference between the N200 response for self- and externally-generated voices.

| **Variable** | **Estimate** | **SE** | **t value** | **Pr(>\|t\|)** |
|---|---|---|---|---|
| **Fixed Effects** | | | | |
| Intercept | -1.160e+00 | 1.900e-01 | -6.104 | 1.28e-06 *** |
| AO | 1.717e-01 | 5.411e-02 | 3.173 | 0.00172 ** |

| | | | | |
|---|---|---|---|---|
| LSHS AVH | 9.422e-04 | 5.226e-02 | 0.018 | 0.98575 |
| 60N | 9.656e-02 | 6.312e-02 | 1.530 | 0.12744 |
| 50N | 1.458e-01 | 6.312e-02 | 2.311 | 0.02175 * |
| 40N | 1.741e-01 | 6.312e-02 | 2.759 | 0.00628 ** |
| Pleasure | 1.286e-01 | 6.312e-02 | 2.037 | 0.04278 * |
| AO*LSHS AVH | 1.033e-02 | 1.522e-02 | 0.679 | 0.49802 |
| **Groups** | **Name** | **Variance** | **SD** | |
| **Random Effects** | | | | |
| Subjects | Intercept | 0.44989 | 0.6707 | |
| Residual | | 0.09959 | 0.3156 | |
| Number of observations: 250, Subjects: 25 | | | | |

### Figure and figure legends

**Supplementary figure 1:** Post experiment stimuli rating. A) Arousal rating on a scale of 1-9 for each voice stimulus. B) Valence rating on a scale of 1-9 for each voice stimulus. C) Ownness rating on a scale of 1-10 for each voice stimulus.

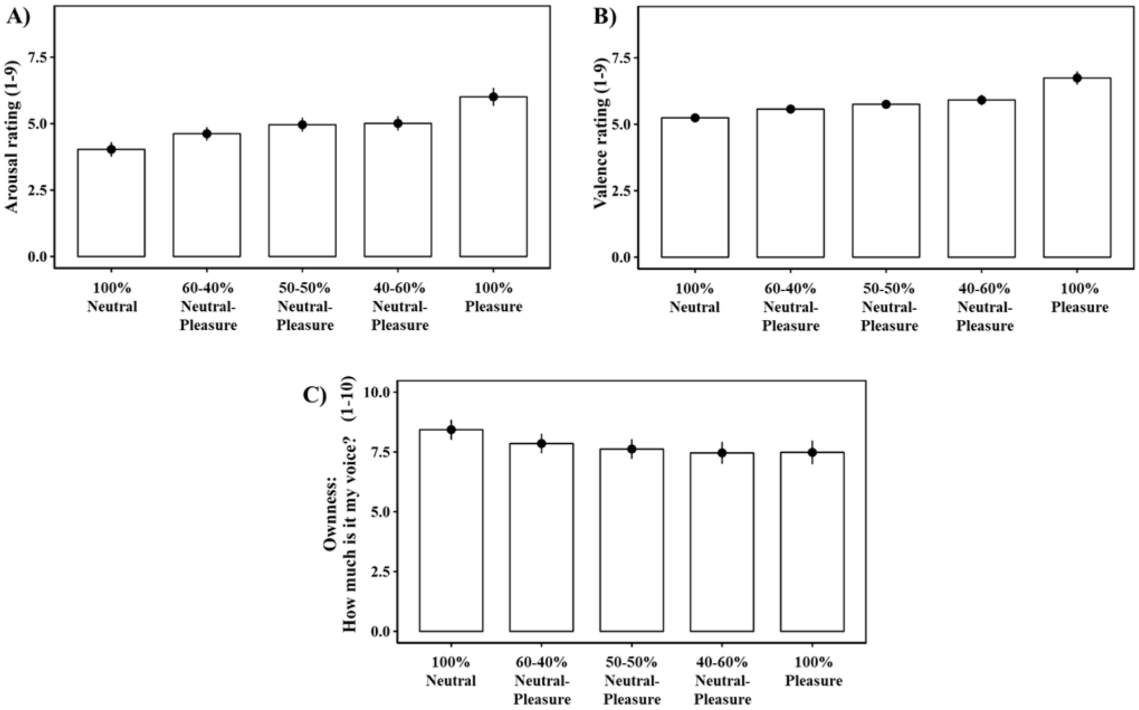

**Supplementary figure 2:** Mean ERP amplitudes for MAc and AO, and suppression effects (AO - MAc) per stimulus type. Note: Negative N100 suppression values depict AO > MAc whereas positive N100 suppression values depict MAc > AO

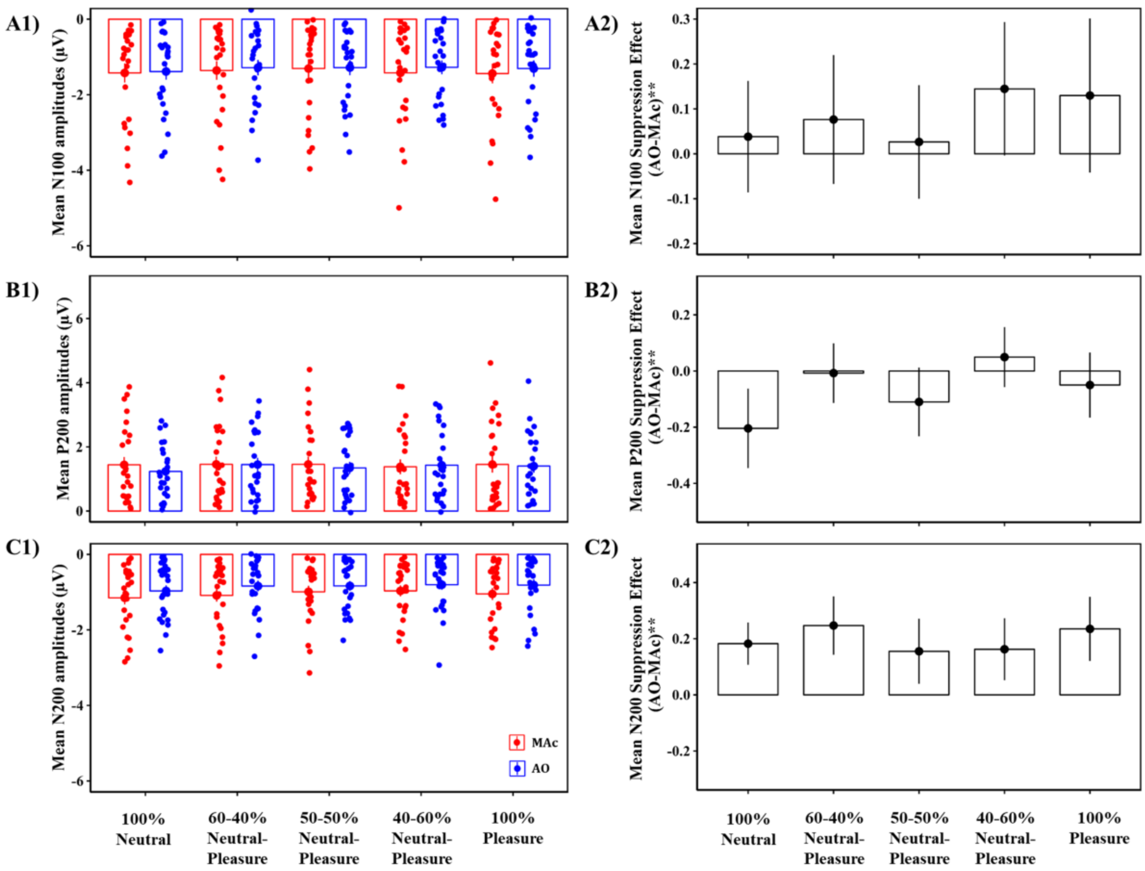

**Supplementary figure 3:** Scatter plots depicting the change in N100 amplitudes as a function of HP (based on LSHS total scores) for each stimulus type.

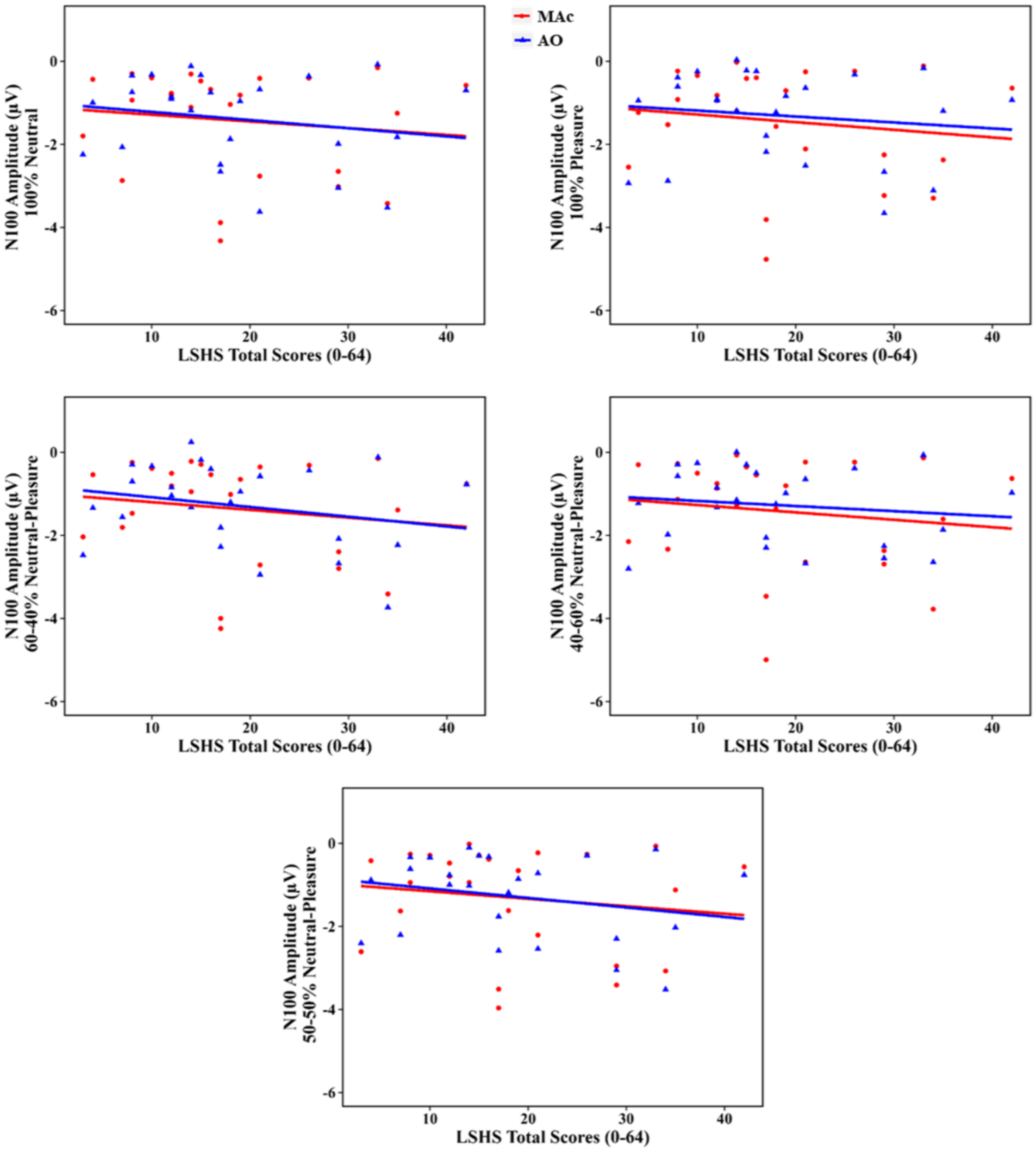

**Supplementary figure 4:** Scatter plots depicting the change in N100 amplitudes as a function of HP (based on LSHS AVH scores) for each stimulus type.

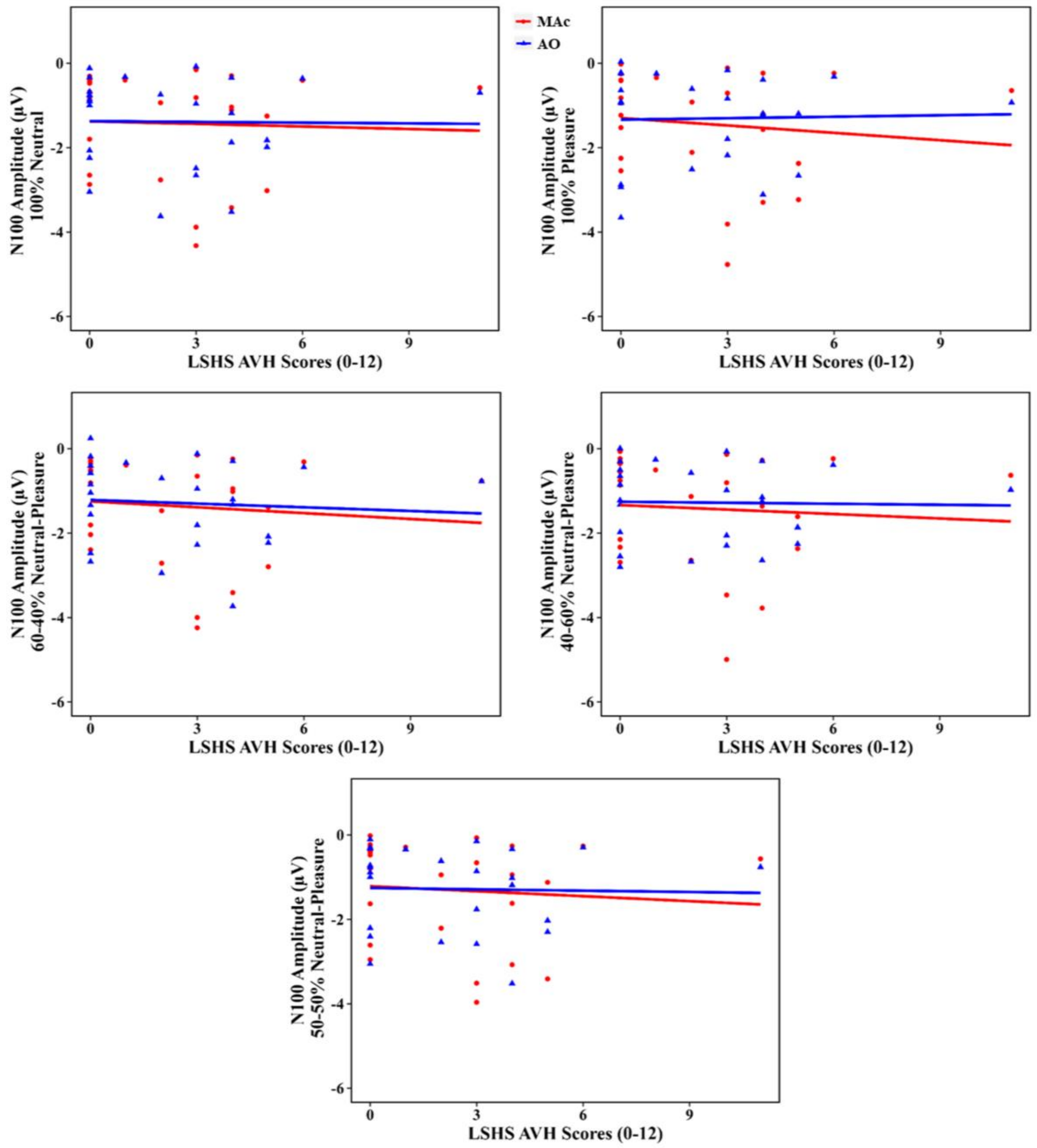

**Supplementary figure 5:** Scatter plots depicting the change in P200 amplitudes as a function of HP (based on LSHS total scores) for each stimulus type.

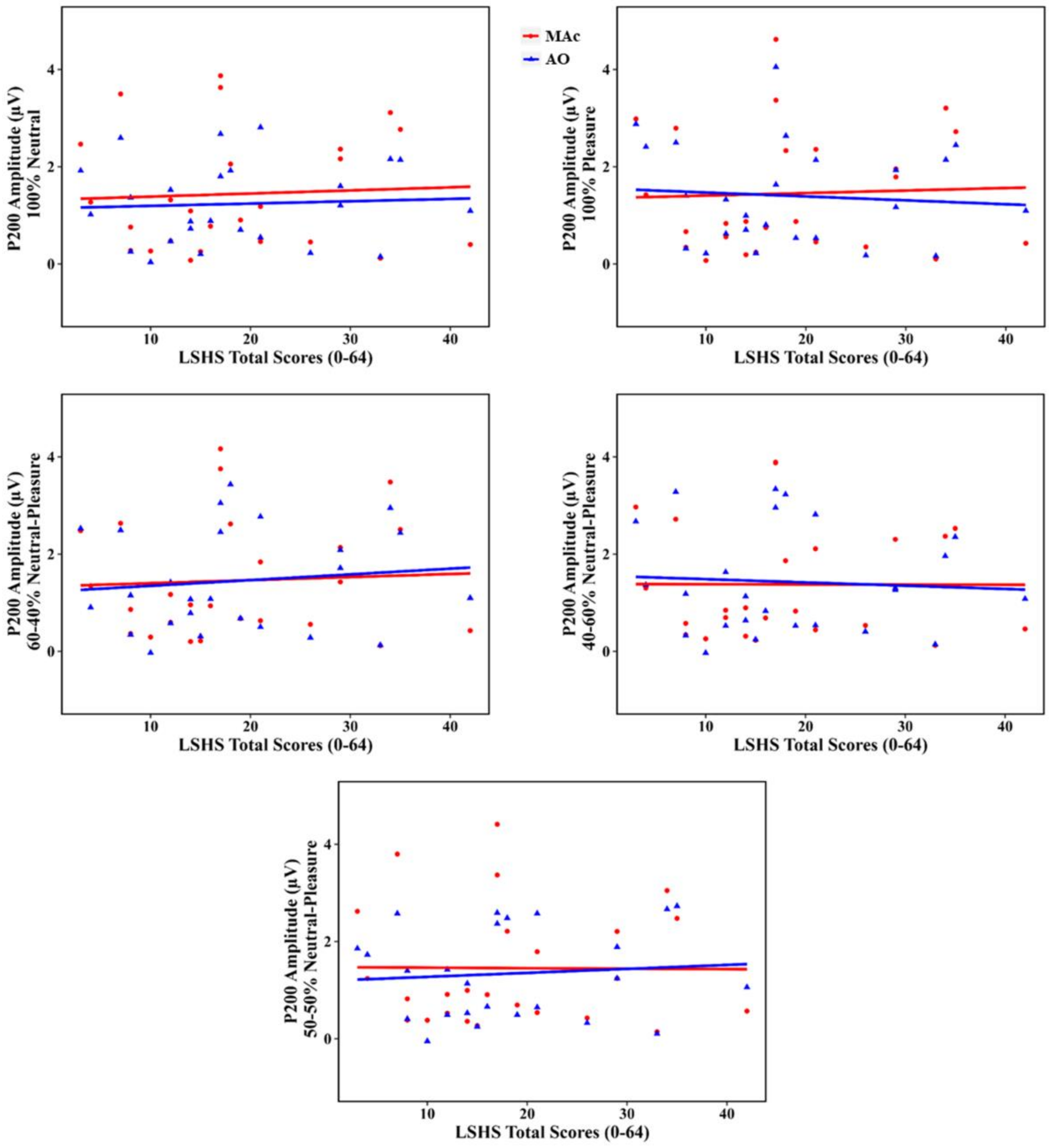

**Supplementary figure 6:** Scatter plots depicting the change in P200 amplitudes as a function of HP (based on LSHS AVH scores) for each stimulus type.

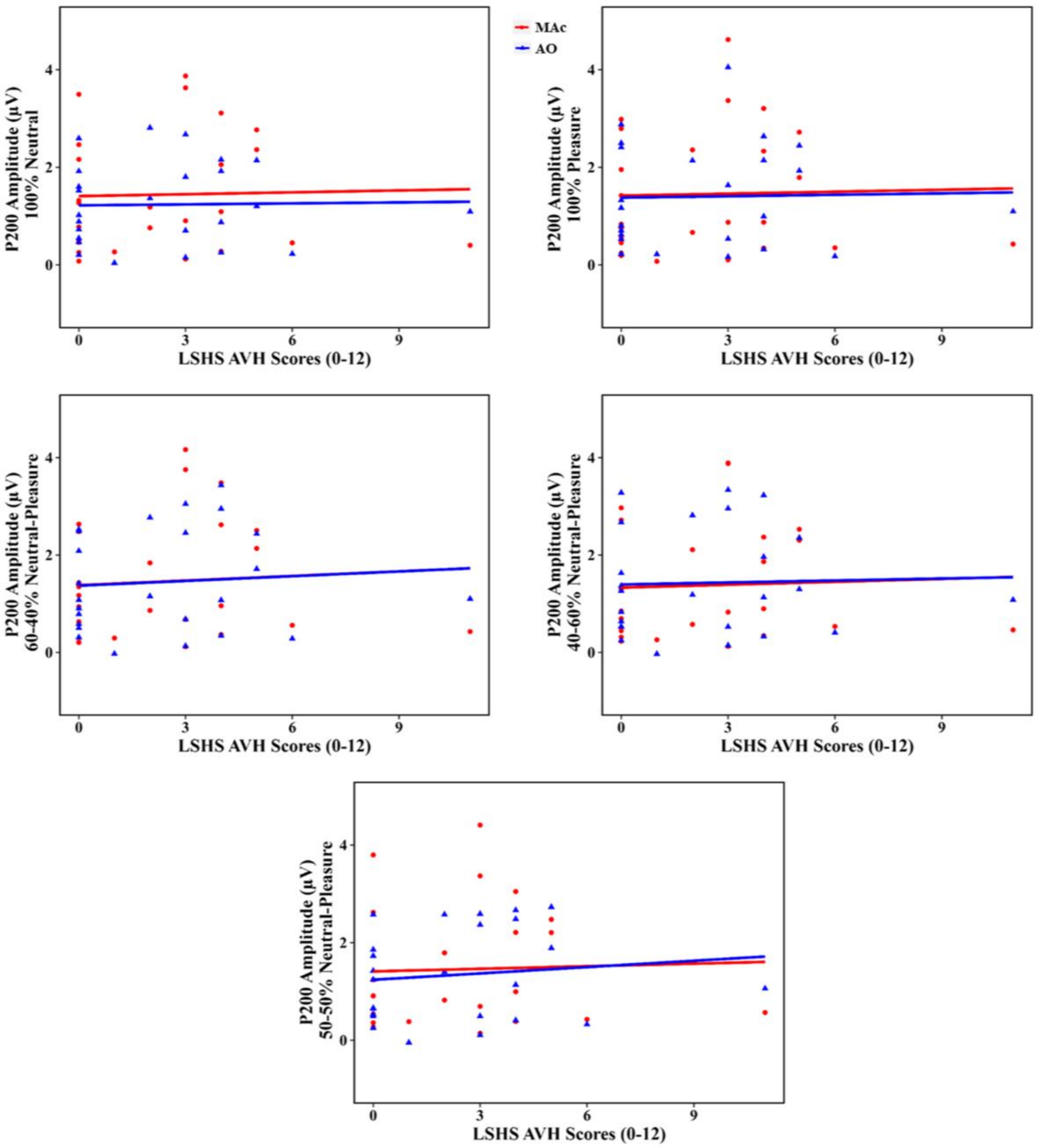

**Supplementary figure 7:** Scatter plots depicting the change in N200 amplitudes as a function of HP based on LSHS AVH scores for each stimulus type. Regardless of the voice quality, the N200 response for the self- and externally-generated voices were significantly different. However, HP did not modulate the N200 response from self- and externally-generated voices.

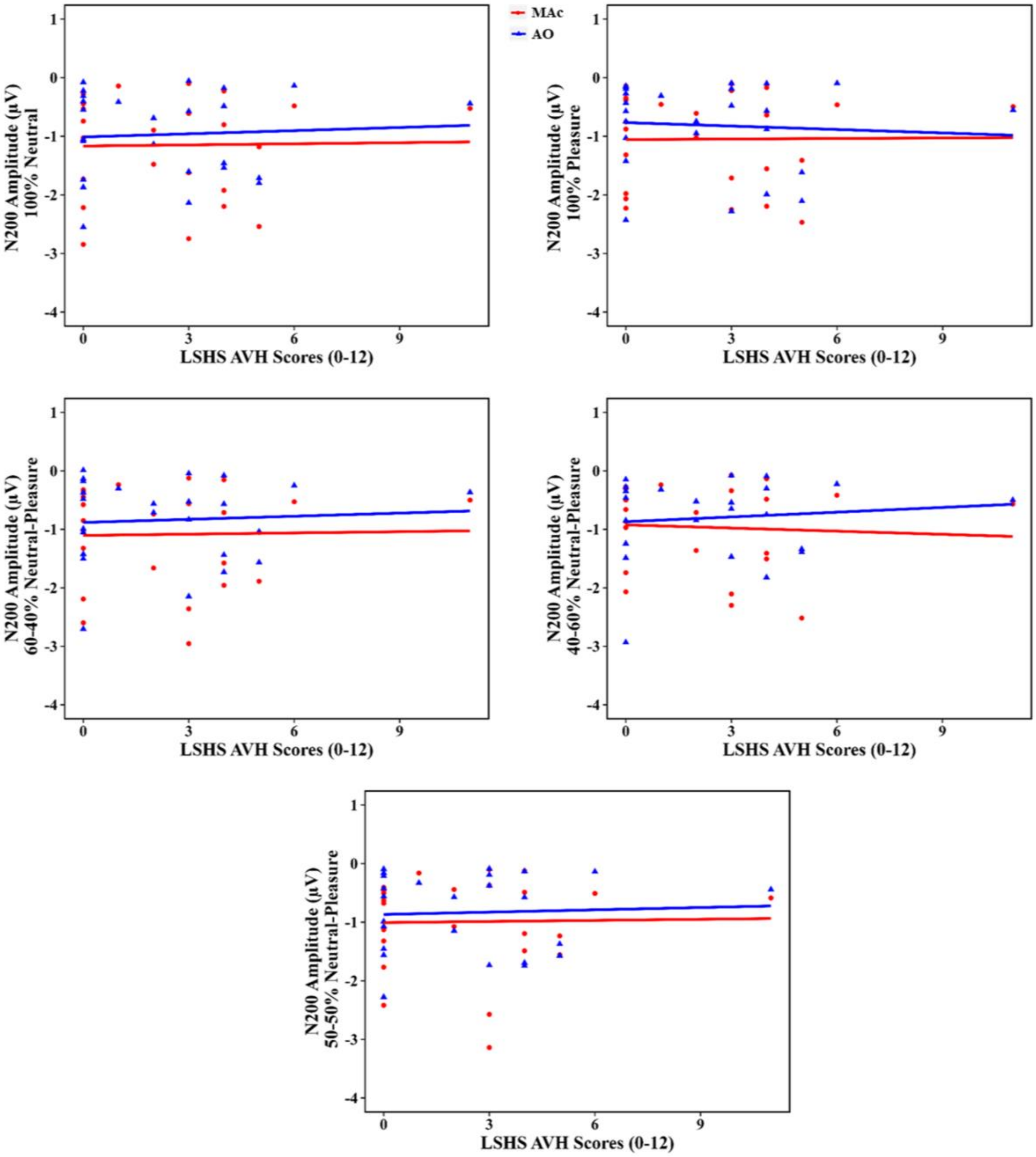
